# *In vitro* and *in vivo* activity of Gepotidacin against drug-resistant mycobacterial infections

**DOI:** 10.1101/2022.08.19.504542

**Authors:** Mohammad Naiyaz Ahmad, Tanu Garg, Shriya Singh, Richa Shukla, VK Ramya, Parvinder Kaur, Sidharth Chopra, Arunava Dasgupta

**Affiliations:** Division of Microbiology, CSIR-Central Drug Research Institute, Sitapur Road, Sector 10, Janakipuram Extension, Lucknow-226031, Uttar Pradesh, India; Academy of Scientific and Innovative Research (AcSIR), Ghaziabad 201002, India; Foundation for Neglected Diseases Research (FNDR), Plot 20A, KIADB Industrial Area, Veerapura, Doddaballapur, Bangalore – 561203, Karnataka, India

**Keywords:** Gepotidacin, *Mycobacterium fortuitum*, *Mycobacterium abscessus*, *Mycobacterium tuberculosis*, drug-resistance, DNA gyrase, Fluoroquinolones, Non tuberculous mycobacteria

## Abstract

Mycobacterial pathogens including Non-tuberculous mycobacteria (NTM) and *M. tuberculosis* (Mtb), are pathogens of significant worldwide interest owing to inherent drug resistance to a wide variety of FDA-approved drugs as well as causing a broad range of serious infections. Identifying new antibiotics active against mycobacterial pathogens is an urgent unmet need, especially those antibiotics that can bypass existing resistance mechanisms. In this study, we demonstrate that Gepotidacin, a first-in-class triazaacenapthylene topoisomerase inhibitor, shows potent activity against Mtb and *M. fortuitum* as well as against other NTMs species, including fluoroquinolone-resistant *M. abscessus.* Furthermore, Gepotidacin exhibits concentration-dependent bactericidal activity against various mycobacterial pathogens, synergizes with several drugs utilized for their treatment, and reduces bacterial load in macrophages in the intracellular killing assay comparable to amikacin. Additionally, *M. fortuitum* ATCC 6841 was unable to generate resistance to Gepotidacin *in vitro*. When tested in a murine neutropenic *M. fortuitum* infection model, Gepotidacin caused a significant reduction in bacterial load in various organs at 10 fold lower concentration than amikacin. Taken together, Gepotidacin possesses a potentially new mechanism of action that enables it to escape existing resistance mechanisms. Thus, it can be projected as a potent novel lead for the treatment of mycobacterial infections, particularly for NTM, where present therapeutic interventions are very limited.

## Introduction

Mycobacterial pathogens, including *M. tuberculosis* (Mtb), the causative agent of Tuberculosis (TB), and Non-tuberculous mycobacteria (NTM) such as *M. fortuitum* and *M. abscessus*, are some of the most formidable pathogens encountered by modern healthcare systems worldwide (1). This is usually due to their ability to cause complex, chronic infections, which are refractory to chemotherapeutic intervention. The discovery and development of new drugs acting against drug-resistant mycobacterial pathogens are an urgent, unmet critical need (2).

Fluoroquinolones (FQ) target DNA gyrase, a clinically validated, broad-spectrum drug target (3). Additionally, FQ are extensively utilized as front-line therapy for treatment of NTM infections, while they are the second line of treatment for infections caused due to Multi-drug resistant (MDR)-TB (4). Indeed, Moxifloxacin (MXF), a fourth-generation FQ, is under multiple clinical trials as a part of combination therapy to reduce the time duration of anti-mycobacterial treatment (5). However, due to extensive clinical utilization, ever-increasing resistance is being clinically encountered, thus nullifying their advantages (6). Hence, identifying and developing novel chemical entities acting against DNA gyrase would be a welcome addition to the anti-bacterial chemotherapeutic arsenal.

In this context, Gepotidacin (GEPO, GSK21409440) is a novel, first-in-class triazaacenapthylene topoisomerase inhibitor with a molecular weight of 448.5, selectively inhibits bacterial type II topoisomerases by interacting with GyrA subunit of bacterial DNA gyrase and ParC subunit of bacterial topoisomerase IV through a unique mechanism that is not utilized by any antibiotic currently in clinical practice (7, 8). Due to this novel mechanism, GEPO is potently active against all bacterial strains, including FQ-resistant strains (7). This is because GEPO’s binding site is close to but distinct from FQ, thus allowing for its activity against FQ-resistant strains (8). Clinical safety of GEPO has been evaluated in healthy human subjects and subjects with renal impairment in phase I clinical trials (NCT02202187, NCT02729038) by both intravenous and oral formulations and has been found to be safe (9, 10, 11, 12). It is currently under various clinical trials for the treatment of sexually transmitted infections (STI) caused by *N. gonorrhoeae* as well as acute bacterial skin and skin structure infections (ABSSSI) (13, 14). However, its activity against mycobacterial pathogens, including NTM, has not been explored yet. Here, in this study, we demonstrate the promising activity of GEPO against various clinically relevant mycobacterial pathogens, including against FQ-resistant strains.

## Materials and Methods

### Media and chemicals

All bacterial media and supplements, including Middlebrook 7H9 broth, Middlebrook 7H11 agar, ADC (albumin, dextrose, and catalase), and OADC (oleic acid, albumin, dextrose, and catalase) supplements, were purchased from BD (Franklin Lakes, NJ, USA). All other chemicals and antibiotics were procured from Sigma-Aldrich (St. Louis, MO, USA). GEPO was purchased from Medchem Express, NJ, USA.

### Bacterial cultures

The bacterial strains including drug-susceptible Mtb H37RvATCC 27294, Isoniazid (INH) resistant Mtb ATCC 35822, Rifampicin (RIF) resistant ATCC 35838, Streptomycin (STR) resistant ATCC 35820, Ethambutol (ETB) resistant ATCC 35837, *M. fortuitum* ATCC 6841, *M. abscessus* ATCC 19977 and *M. chelonae* ATCC 35752 were procured from ATCC (ATCC, Manassas, USA). The mycobacteria were propagated in Middlebrook 7H9 broth supplemented with ADC and 0.05% tween-80 at 37°C. Middlebrook 7H11 agar supplemented with 0.2% glycerol and 10% OADC plates were used for colony forming unit (CFU) enumeration.

### Antibacterial susceptibility testing

Antibacterial susceptibility testing was carried out utilizing broth microdilution assay according to CLSI guidelines (15). 10 mg/mL stock solutions of test and control compounds were prepared in DMSO and stored at −20°C. Bacterial cultures were inoculated in supplemented Middlebrook 7H9 broth(Middlebrook 7H9 broth supplemented with 10% ADC, 0.2% glycerol, 0.05% tween-80) with a density of ~10^6^ CFU/mL based on OD_600_. The compounds were tested from 64-0.5 mg/L in a two-fold serial diluted fashion with 2.5 μL of each concentration added per well of a 96-well round bottom microtiter plate. Later, 97.5 μL of bacterial suspension was added to each well-containing test compound along with controls and incubated at 37°C for 7 days for slow-growing mycobacteria and 48 h for fast-growing NTM. MIC is defined as the lowest compound concentration where there is no visible growth. MIC assays were performed in duplicates, and each assay was carried out independently three times.

### Bacterial time kill kinetics

Bactericidal activity of GEPO was assessed by the time-kill method (16). *M. fortuitum* ATCC 6841, *M. abscessus* ATCC 19977, and Mtb H37Rv ATCC 27294 were diluted to achieve ~10^6^ CFU/mL in a total volume of ~0.3 mL and added to 96 well plates along with GEPO and controls at 1x, and 10x MIC followed by incubation at 37°C for 7 days for slow-growing mycobacteria and 48 h for NTM. For evaluating the reduction in CFU, 0.03 mL sample was removed at various time-points, serially two-fold diluted in 0.27 mL normal saline, and 0.1 mL of respective dilution was spread on supplemented Middlebrook 7H11 agar plate (Middlebrook 7H11 agar supplemented with10% OADC and0.2% glycerol). The plates were incubated at 37°C for 72 h for NTM, 4 weeks for slow-growing mycobacteria, and colonies were enumerated. Kill curves were constructed by counting the colonies from plates and plotting CFU/mL of surviving bacteria at each time point in the presence and absence of the compound. Each experiment was performed in duplicates and repeated three times independently, and the mean data was plotted.

### Synergy with front-line anti-mycobacterial drugs

Determination of interaction of GEPO with typically utilized antibiotics for mycobacterial treatment, including Amikacin (AMK), Meropenem (MEM), Moxifloxacin (MXF), Ciprofloxacin (CIP), Linezolid (LZD), Clarithromycin (CLR), and Vancomycin (VAN) was tested by checkerboard method as per CLSI guidelines. Serial two-fold dilutions of each drug were freshly prepared prior to testing. GEPO was two-fold diluted along the ordinate (8 dilutions) while antibiotics were serially diluted along the abscissa (12 dilutions) in 96 well microtiter plate. 95 μL of ~10^5^ CFU/mL of Mtb, *M. fortuitum* ATCC 6841, and *M. abscessus* ATCC 19977 was added to each well, and plates were incubated at 37°C for 48 h for NTM and 7 days for slow-growing mycobacteria. After the incubation period was over, ΣFICs (fractional inhibitory concentrations) were calculated as follows: ΣFIC = FIC A + FIC B, where FIC A is the MIC of drug A in the combination/MIC of drug A alone and FIC B is the MIC of drug B in the combination/MIC of drug B alone. The combination is considered synergistic when the ∑FIC is ≤0.5, additive or indifferent when the ∑FIC is >0.5 to 4, and antagonistic when the ∑FIC is >4 (17).

### Activity of GEPO against intracellular mycobacteria

*M. fortuitum* ATCC 6841 was grown overnight in supplemented 7H9 Middlebrook broth. The bacterial inoculum was prepared from mid-log phase bacteria and diluted to ~10^7^cfu/mL. ~10^5^ cells/well of J774A.1 cells were seeded in 6-well flat bottom plates and infected with *M. fortuitum* at 1:5 MOI. After 4 hours of infection, cells were washed twice with 1x Phosphate buffer saline (PBS), replenished with fresh RPMI medium containing GEPO and AMK, and further incubated for 48 h. To estimate initial count, 3 wells were lysed 4 h post-infection, serially diluted, and plated on supplemented Middlebrook 7H11 agar plate. After 96 h of incubation, 3 wells were lysed, serially diluted, plated, and incubated at 37°C for 72 h to estimate CFU. Kill curves were constructed by counting the colonies from plates and plotting the CFU/mL. Each experiment was repeated three times in duplicate, and the mean data was plotted.

### Determination of Post antibiotic effect (PAE) of GEPO

To determine PAE of GEPO, *M. fortuitum* ATCC 6841 was diluted in supplemented 7H9 Middlebrook broth to achieve ~10^5^ CFU/mL and exposed to 1x and 10x MIC of GEPO, AMK, and LVX and incubated at 37°C for 1 h. Following the incubation period, the culture was centrifuged and washed 2 times with pre-warmed supplemented 7H9 Middlebrook broth to remove any traces of antibiotics. Finally, cells were re-suspended in drug-free, supplemented 7H9 Middlebrook broth, and incubated further at 37°C. After every 3h, samples were taken serially diluted and plated on supplemented Middlebrook 7H11 agar plate for enumeration of CFU. The PAE was calculated as PAE = T – C; where T is the difference in time required for 1 Log_10_ increase in CFU after drug exposure and C is the time difference for 1 Log_10_ increase in CFU in similarly treated drug-free control (18).

### *In vitro* resistant mutants generation and resistant frequency

*In vitro* emergence of GEPO resistance and determination of resistance frequency were performed as reported (19, 20). ~10^8^ CFU of *M. fortuitum* was spread plated on supplemented Middlebrook 7H11 agar plates containing 8-256x MIC of GEPO, CLR, and LVX along with supplemented Middlebrook 7H11 agar plates without antibiotics and incubated at 37°C, whereas, for calculating mutant prevention concentration (MPC), ~10^10^ CFU of *M. fortuitum* was spread plated on drug-containing plates (31). The plates were observed daily from the 3^rd^-14^th^ day for the appearance of colonies. The resistant colonies from these plates were streaked over fresh drug-containing plates and the days taken to form colonies on fresh drug plates were compared with the day of appearance of that particular colony in the previous step. The resistant frequency (R_f_) is calculated as R_f_ = R_N_/T_N,_ where R_N_ is the number of resistant colonies on the drug plate, and T_N_ is the CFU of inoculum (*i.e.*, CFU on the drug-free plate x Dilution factor). A similar experiment was also performed on 14^th^-day old drug-containing plates stored at 37°C to compare mutant generation frequency on fresh drug-containing supplemented 7H11 agar plates with 14^th^-day old plates to rule out the emergence of resistance due to stability issues or possible degradation of the drug. Furthermore, we transferred individual resistant mutant colonies to drug-free 7H9 broth and grew them to mid-log phase (OD_600_- 0.4-0.8), then subsequently given 5 passages on drug-free 7H9 broth. Afterward, the MIC of different antibiotics against these *in vitro* resistance mutants was determined as described above.

### Animal experiments

Animal experiments were performed on six-eight week old BALB/c mice procured from the National Laboratory Animal Facility of CSIR-Central Drug Research Institute, Lucknow. The Institutional Animal Ethics Committee approved the experimental protocols.

### *In vivo* evaluation of GEPO in murine neutropenic bacteremia model of *M. fortuitum*

Since the majority of NTM infections occur under immuno-compromised conditions, we used a murine neutropenic *M. fortuitum* infection model to mimic clinical conditions as described previously (21). Briefly, six-week-old (22-24 g) male BALB/c mice were randomly divided into 4 groups: untreated control (nine mice, Group 1), GEPO treated (six mice, Group 2), LVX treated (six mice, Group 3) and AMK treated (six mice, Group 4). Neutropenia was induced in mice by IP doses of cyclophosphamide 150 mg/Kg, administered 4 days and 1 day before infection. Infection in mice was initiated by an IV injection of ~5×10^6^ CFU *M. fortuitum* ATCC 6841 in the tail vein. The infected mice were treated daily with a single dose of GEPO (orally, 10 mg/Kg in 5% acacia gum solution) in Group 2, LVX (orally, 10 mg/kg in 5% acacia gum solution) in Group 3 and AMK (IP, 100 mg/Kg in PBS) in Group 4 for fifteen days, while control mice in Group 1 were given with placebo (oral, 5% acacia gum solution in water, and/or PBS through IP). On the eighteenth day post-infection, all mice were sacrificed, various organs were resected, homogenized and bacterial burden was enumerated by plating on supplemented 7H11 Middlebrook agar plates. Kill curves were constructed by counting the colonies from plates and plotting the CFU/mL. Each experiment was done in duplicate, and CFU of individual mice was plotted.

### Statistical analysis

Statistical analysis was performed using GraphPad Prism 6.0 software (GraphPad Software, La Jolla, CA, USA). Comparison between three or more groups was analyzed using one-way ANOVA, with post-hoc Tukey’s multiple comparisons test. P-values of <0.05 were considered to be significant.

## Results and Discussion

### *In vitro* anti-mycobacterial activity

The MIC of GEPO was determined against a mycobacterial strain panel including Mtb, and various clinically important NTM’s and MIC results are tabulated in Table 1 (A and B). As can be seen in Table 1A, GEPO exhibits significantly lower MIC (2 mg/L) against fast-growing NTM such as *M. fortuitum* ATCC 6841*, M. chelonae* ATCC 35752,andFQ-resistant *M. abscessus* ATCC 19977 and its MIC is comparable to AMK, which is one of potent therapeutic intervention against NTM infections (22). However, MIC of GEPO was higher than fluoroquinolones (MXF, LVX, CIP) against most mycobacterial pathogens; additional structural modification and optimization may enhance its activity against mycobacteria. Interestingly, MIC of GEPOagainst tested mycobacterial pathogens is in line with reported MIC of GEPO against various other bacterial pathogens (≤0.06 mg/L to 2 mg/L)for which GEPO is currently being evaluated in various phases of clinical trials (25). However, GEPO is much less potent against slow-growing NTM such as *M. avium* ATCC 19698, *M. gordonae* ATCC 14470, *M. kansasii* ATCC 12478, *M. intracellulare* ATCC 13950 and *M. nonchromogenicum* ATCC 19530 (MIC 16-32 mg/L). At the same time, AMK is almost equipotent across NTM except for *M. abscessus* ATCC 19977, where its activity is inferior to GEPO (MIC 8 mg/L). This is significant because *M. abscessus* ATCC 19977 is one of the most drug-resistant mycobacterial pathogen (23, 24). When tested for *in vitro* anti-tuberculosis activity against Mtb H37Rv ATCC 27294, GEPO showed very promising activity (MIC 0.5 mg/L) and equipotent activity against clinical drug-resistant Mtb strains (Table 1B). The MIC of GEPO against various mycobacterial strains compares very well with reported MIC against various bacterial pathogens, including those resistant to FQs (25). This equi-potency against multiple clinical mycobacterial pathogens demonstrates that GEPO exhibits a new mechanism of action that is not influenced by existing drug-resistance mechanisms.

**Table 1:**
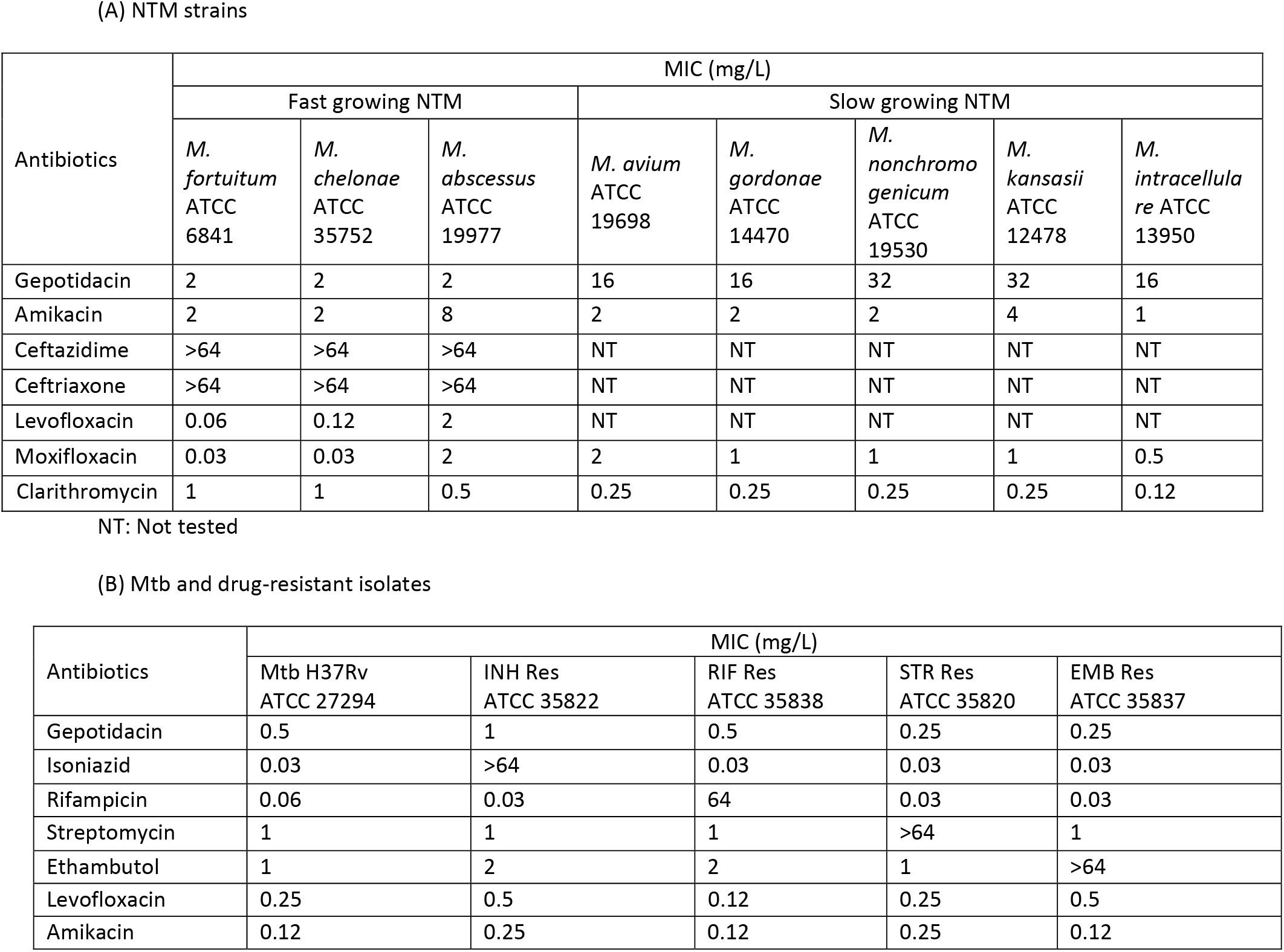
MIC (mg/L) of GEPO and other comparator drugs against various mycobacterial strains (A) NTM strains and (B) Mtb resistant isolates

### *In vitro* bactericidal activity determination by time-kill kinetic assay

To determine the killing kinetics of GEPO against various mycobacteria, GEPO was tested at 1× and 10× MIC against NTM’s and Mtb along with control drugs, and kill curves were plotted in Fig 1A-C. As shown in Fig 1A-C, GEPO exhibits concentration-dependent bactericidal activity against all three mycobacterial strains tested. When tested against *M. fortuitum* ATCC 6841, 1x MIC of GEPO caused a reduction of ~5 log_10_ CFU/mL. In contrast, 10x MIC of GEPO eliminated culture (~9 log_10_ CFU/mL reduction) in 48 h with no observed re-growth compared to untreated control (Fig 1A). This concentration-dependent killing is comparable to LVX, which caused a reduction of ~8.5 log_10_ CFU/mL in 24 h at 10x MIC, while its 1x MIC caused a reduction of ~5.2 log_10_ CFU/mL in 24h. In contrast, FQ-resistant *M. abscessus* ATCC 19977 is far less susceptible to LVX (MIC 2 mg/L), with 10x of LVX taking 48 h to reduce ~8.5 log_10_CFU/mL. In contrast, the killing kinetics of GEPO remain undisturbed in the presence of FQ resistance (Fig 1B). This data indicates that the kill kinetics of GEPO is not affected by the FQ susceptibility of the strain.

**Figure 1:**
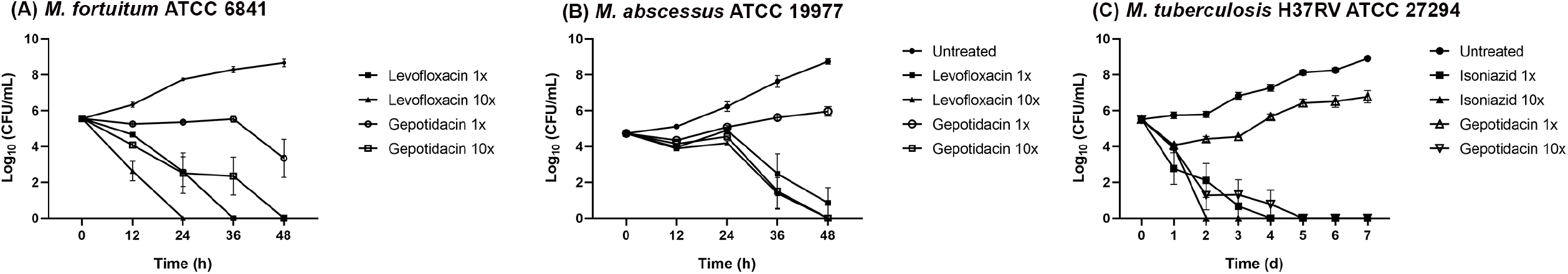
Time kill kinetics of Gepotidacin and comparator antibiotics against various mycobacterial strains (A) *M. fortuitum* ATCC 6841 (B) *M. abscessus* ATCC 19977 & (C) *M. tuberculosis* H37Rv ATCC 27294

Following the same trend, GEPO exhibits concentration-dependent bactericidal activity against Mtb H37Rv ATCC 27294. 10x MIC of GEPO caused a reduction of ~8.0 log_10_ CFU/mL in 120 h and no re-growth afterward, whereas 1x MIC of GEPO led to a reduction of only ~1.7 log_10_CFU/mL in 120 h as compared to untreated control (Fig 1C). Similar concentration-dependent killing kinetics is observed in INH, where 1x MIC and 10x MIC of INH lead to a ~3.6 log10 CFU/mL reduction and complete elimination of culture (reduction of ~5.7 log_10_CFU/mL) respectively in 24 h as can be seen in Fig 1C. Taken together, GEPO exhibits concentration-dependent killing kinetics, which is similar to LVX for NTM’s and INH for Mtb.

### Drug combination assays

The treatment of complex, recalcitrant mycobacterial infections often utilizes various drug combinations, which help in suppressing the emergence of drug resistance as well as lead to a faster clearance of infection, thus improving patient compliance while simultaneously reducing drug-related toxicity (26, 27). This situation demands that any new drug candidate be tested for its ability to synergize with other approved drugs. The ability of GEPO to interact and/or synergize with drugs utilized for the treatment of mycobacterial pathogens was determined by the chequerboard method, which facilitated the calculation of FIC. As can be seen in Table 2, GEPO synergizes with MEM, MXF, CIP, and LZD, whereas it has no interaction with VAN and AMK against *M. fortuitum* ATCC 6841. On the other hand, GEPO synergized with MXF, CIP and AMK against *M. abscessus* ATCC 19977. Interestingly, GEPO synergized with INH against Mtb, whereas it did not interact with RIF, ETB, and STR.

**Table 2:**
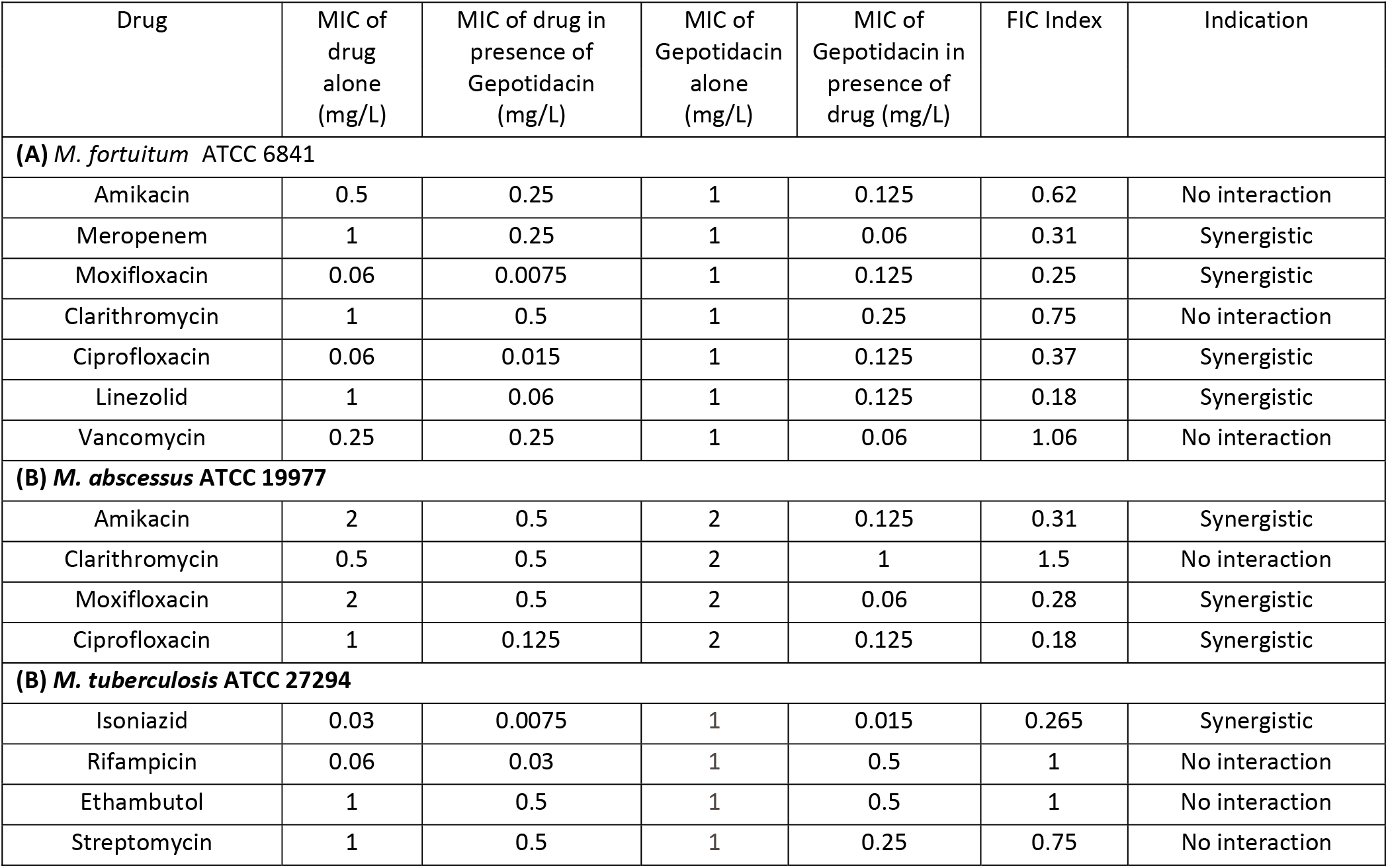
Determination of interaction of GEPO with front line anti-mycobacterial drugs against (A) *M. fortuitum* ATCC 6841, (B) *M. abscessus* ATCC 19977 and (C) *M. tuberculosis* ATCC 27294

| Drug | MIC of drug alone (mg/L) | MIC of drug in presence of Gepotidacin (mg/L) | MIC of Gepotidacin alone (mg/L) | MIC of Gepotidacin in presence of drug (mg/L) | FIC Index | Indication |
| --- | --- | --- | --- | --- | --- | --- |
| **(A) *M. fortuitum* ATCC 6841** |  |  |  |  |  |  |
| Amikacin | 0.5 | 0.25 | 1 | 0.125 | 0.62 | No interaction |
| Meropenem | 1 | 0.25 | 1 | 0.06 | 0.31 | Synergistic |
| Moxifloxacin | 0.06 | 0.0075 | 1 | 0.125 | 0.25 | Synergistic |
| Clarithromycin | 1 | 0.5 | 1 | 0.25 | 0.75 | No interaction |
| Ciprofloxacin | 0.06 | 0.015 | 1 | 0.125 | 0.37 | Synergistic |
| Linezolid | 1 | 0.06 | 1 | 0.125 | 0.18 | Synergistic |
| Vancomycin | 0.25 | 0.25 | 1 | 0.06 | 1.06 | No interaction |
| **(B) *M. abscessus* ATCC 19977** |  |  |  |  |  |  |
| Amikacin | 2 | 0.5 | 2 | 0.125 | 0.31 | Synergistic |
| Clarithromycin | 0.5 | 0.5 | 2 | 1 | 1.5 | No interaction |
| Moxifloxacin | 2 | 0.5 | 2 | 0.06 | 0.28 | Synergistic |
| Ciprofloxacin | 1 | 0.125 | 2 | 0.125 | 0.18 | Synergistic |
| **(C) *M. tuberculosis* ATCC 27294** |  |  |  |  |  |  |
| Isoniazid | 0.03 | 0.0075 | 1 | 0.015 | 0.265 | Synergistic |
| Rifampicin | 0.06 | 0.03 | 1 | 0.5 | 1 | No interaction |
| Ethambutol | 1 | 0.5 | 1 | 0.5 | 1 | No interaction |
| Streptomycin | 1 | 0.5 | 1 | 0.25 | 0.75 | No interaction |

This observed synergy was further tested by combination time-kill analysis, and the results are plotted in Fig2A-C. As shown in Fig 2A, a combination of 1x MIC of LZD and GEPO was dramatically more effective than either LZD and GEPO alone and reduced ~7.5 log_10_ CFU/mL of *M. fortuitum* ATCC 6841 in 48 h. At the same time, the combination of 1x MIC of GEPO and MXF is indistinguishable from 1x MXF alone and eliminates all culture in 24 h compared to untreated. The combination of 1x MIC of GEPO and MEM was more effective than 1x MIC of MEM alone, but it could not be distinguished from killing by 1x MIC of GEPO and both of them reduced ~4.5 log_10_CFU/mL of *M. fortuitum* ATCC 6841 in 48 h.

**Figure 2:**
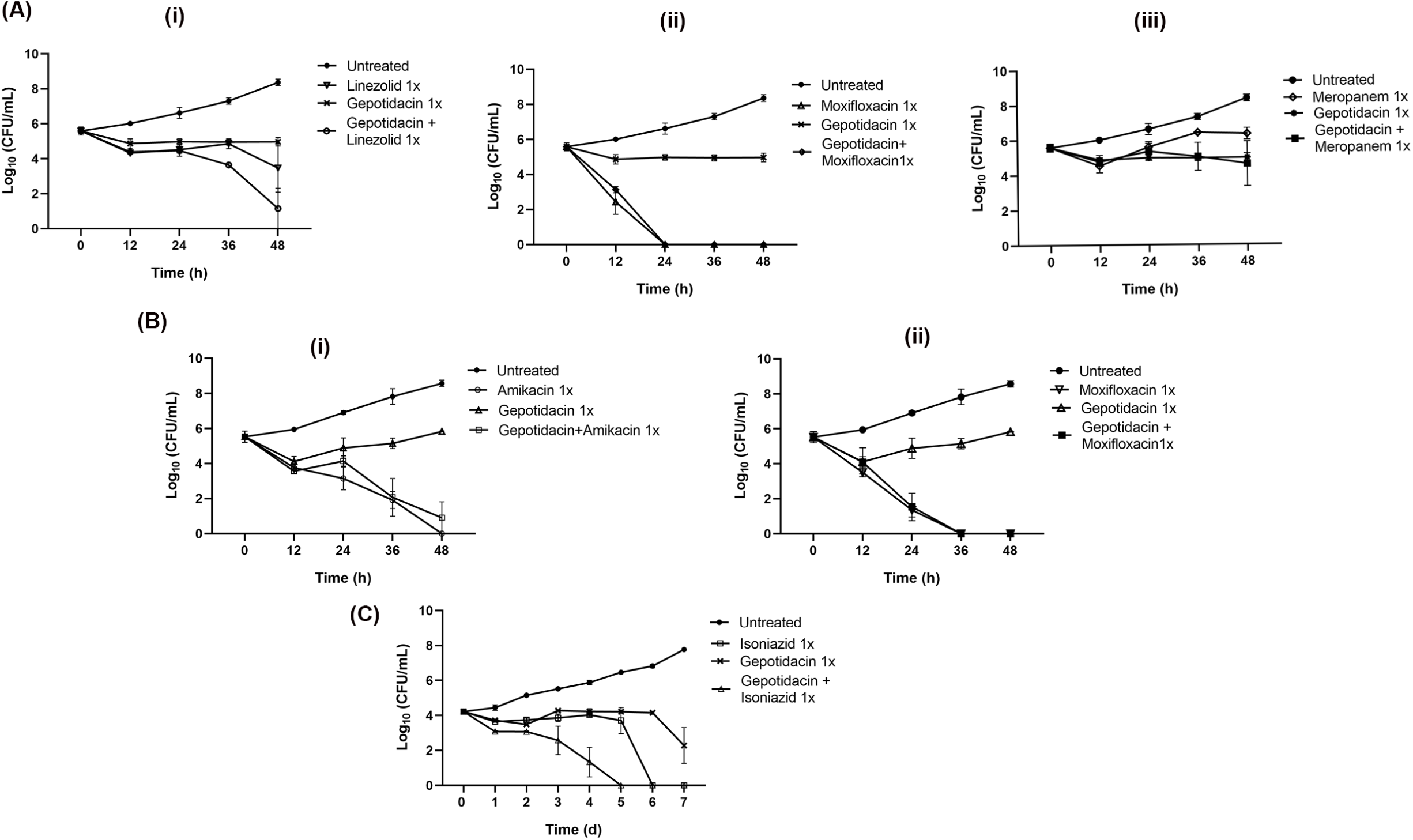
Time kill kinetics of Gepotidacin in combination with (A) (i) LZD, (ii) MXF and (iii) MEM against *M. fortuitum* ATCC 6841, with (B) (i) AMK and (ii) MXF against *M. abscessus* ATCC 19977 and with (C) INH against *M. tuberculosis* ATCC 27294

The combination of 1x MIC of AMK and GEPO was much more potent than 1x MIC of GEPO alone and reduced ~7.5 log_10_ CFU/mL in 48 h for FQ-resistant *M. abscessus* ATCC 19977. The combination efficacy of 1x MIC of GEPO and MXF was indistinguishable from 1x MIC of MXF alone and eliminated culture in 36 h while 1x GEPO alone did not (Fig 2B).

In the case of Mtb, a combination of 1x MIC of INH and GEPO was more effective than either INH and GEPO alone and eliminated all culture (~6.5 log_10_ CFU/mL) in 120 h which is faster than INH (144h), with 1x MIC GEPO alone reducing ~5.5 log_10_CFU/mL in 168 h. No re-growth was observed for any mycobacterial pathogen under any conditions. Collectively, the studies indicate that identified combinations could potentially be utilized as a part of multi-drug therapy to treat drug-resistant recalcitrant mycobacterial infections (Fig 2C).

### Intracellular killing of *M. fortuitum* ATCC 6841

Since mycobacteria are significant intracellular pathogens, the ability of GEPO to clear intracellular mycobacterial infection was determined. Briefly, J774A.1 cells were infected with *M. fortuitum* ATCC 6841. GEPO and AMK were added to the culture at multiple MIC’s, followed by cell lysis and plating on supplemented Middlebrook 7H11 agar to determine remaining CFU’s. As can be seen in Fig 3, GEPO at 5x MIC significantly reduced mycobacterial load in macrophages by ~4 log_10_ CFU/mL. In contrast, AMK at 5x MIC reduced it by ~4.5 log_10_ CFU/mL in 96 h compared to untreated control. This demonstrates the potent ability of GEPO to control and eliminate intracellular mycobacterial growth, which is quite comparable to that of AMK, a frontline antimicrobial typically utilized for the treatment of NTM infections, thus boding well for its utilization as a potent anti-mycobacterial therapeutic.

**Figure 3:**
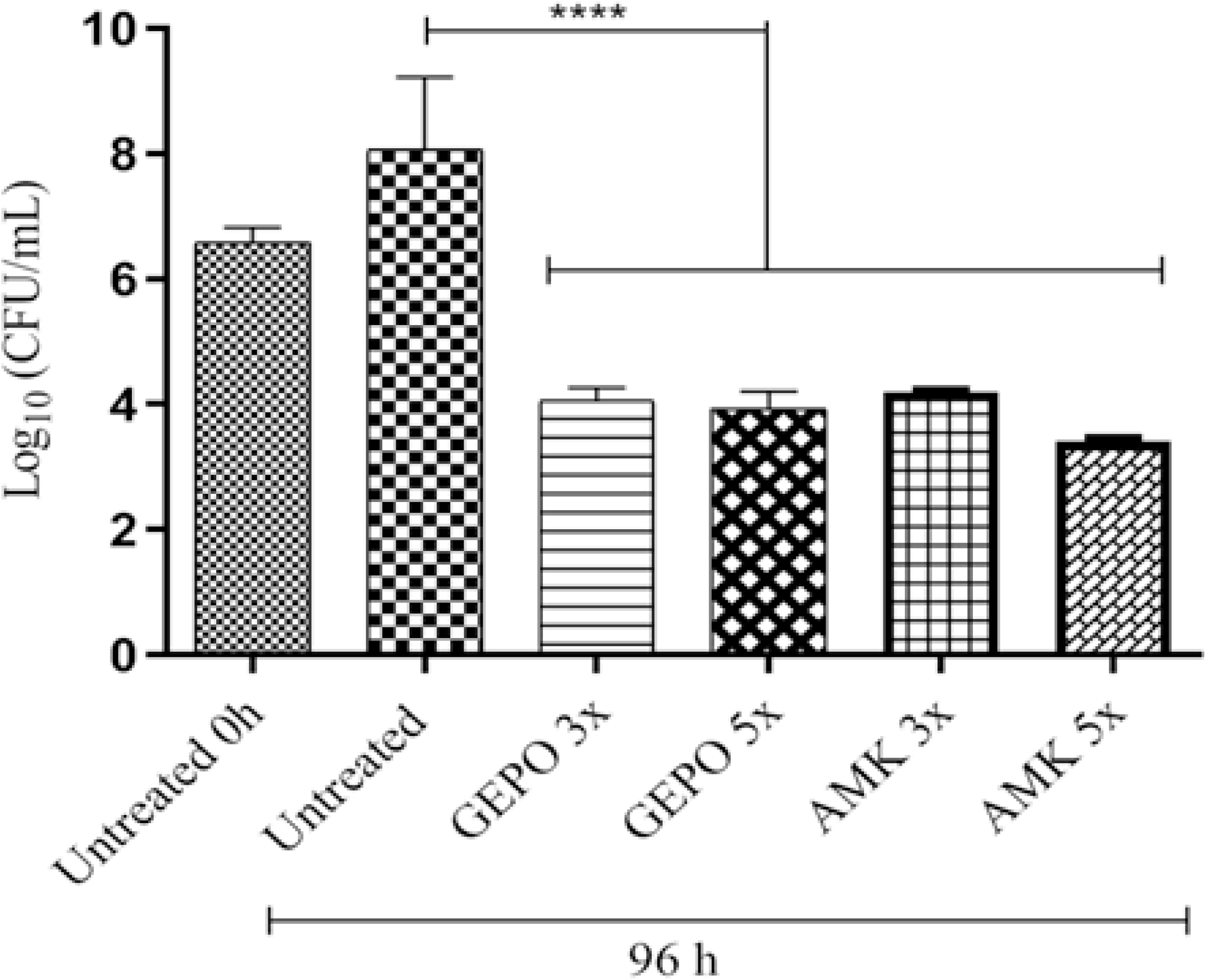
Intracellular activity of Gepotidacin against *M. fortuitum* ATCC 6841 at 96 h. (**** equals <0.0001). All data were presented as mean ± s.d.

### *In vitro* post-antibiotic effect (PAE) against *M. fortuitum* ATCC 6841

Possessing a long PAE is an asset for any antibacterial molecule under consideration as it helps in minimizing the dosages required for therapeutic clearance. In this context, GEPO exhibited a prolonged PAE of 18 h at 10x MIC and 12 h at 1x MIC, which is better than LVX (12 h at 10x MIC, 6 h at 1x MIC) but comparable to AMK (Table 3). This *in vitro* PAE of GEPO against *M. fortuitum* is comparable to earlier reports of ~3-12 h against S*. aureus* under *in vivo* conditions (29) and could be a consequence of GEPO inhibiting dual targets (7, 8). Thus, GEPO exhibits concentration-dependent bactericidal activity with prolonged PAE.

**Table 3:** Determination of Post Antibiotic effect (PAE) of GEPO against *M. fortuitum* ATCC 6841

| Treatments | Time taken for 1 log<sub>10</sub> growth (h) | PAE (h) |
| --- | --- | --- |
| UNTREATED | 12 | 0 |
| Amikacin 1x MIC | 24 | 12 |
| Amikacin 10x MIC | 30 | 18 |
| Levofloxacin 1x MIC | 18 | 6 |
| Levofloxacin 10x MIC | 24 | 12 |
| Gepotidacin 1x MIC | 24 | 12 |
| Gepotidacin 10x MIC | 30 | 18 |

### *In vitro* spontaneous resistant mutant generation and resistance frequency

Considering the unrelenting rise in antimicrobial resistance (AMR), possessing a low resistance frequency is a must-have property for any investigational new anti-infective agent. When tested for its ability to induce resistance in *fortuitum* ATCC 6841, we were unable to isolate any stable resistant mutants to GEPO. In contrast, under the same conditions, *M. fortuitum* ATCC 6841 was able to generate stable resistant mutants to LVX and CLR. The resistance frequency for GEPO was calculated to be ~10^−9^ as per the number of colonies that appeared till 14^th^ day upon inoculation on 7H11-agar plates containing 16x MIC of GEPO, which is substantially lower than the resistance frequency of LVX (~10^−5^) at same concentration. On the other hand, we observed lawn-like growth on CLR-containing plates after the 7-14^th^ days, thus unable to calculate the resistance frequency.

Our observed resistance frequency for LVX against *M. fortuitum* (Table 4) is in line with the previously reported resistance frequency for ciprofloxacin (~10^−5^) against *E. coli* in 5-13 days (19). However, we didn’t observe any changes in the number of spontaneous resistant mutant colonies when we extended incubation time as it was observed by Cirz*et al*., (19), rather than it was dependent on the concentration of the drug (Table 4) most likely due to difference in the method and organisms. Additionally, we performed antimicrobial susceptibility assays for *in vitro* resistant mutant isolates of *M. fortuitum* recovered from LVX and CLR containing agar plates after growing them on drug-free media. We didn’t observe any cross-resistance to GEPO in LVX and CLR-resistant isolates (Table 5), whereas isolates recovered from GEPO-containing agar plates didn’t reproduce resistance against GEPO after subsequent passage on drug-free media.

**Table 4:** Frequency of spontaneous resistance mutants and Mutant prevention concentration (MPC) for Levofloxacin and Gepotidacin against *M. fortuitum* ATCC 6841.

| Drugs | MIC (mg/L) | Mutant Frequency and Fold MIC |  |  |  |  |  | Mutant Prevention Concentration (MPC) |
| --- | --- | --- | --- | --- | --- | --- | --- | --- |
|  |  | 8x | 16x | 32x | 64x | 128x | 256x |  |
| Levofloxacin | 0.06 | $0.8 \times 10^{-5}$ | $1.8 \times 10^{-5}$ | $3.6 \times 10^{-7}$ | $3.5 \times 10^{-7}$ | $2.8 \times 10^{-8}$ | 0 | 256x MIC (15.36 mg/L) |
| Gepotidacin | 1 | $1.1 \times 10^{-7}$ | $3.7 \times 10^{-9}$ | 0 | 0 | 0 | 0 | 32x MIC (32 mg/L) |

**Table 5:** MIC of different antibiotics against *in vitro* spontaneous resistant mutant isolates of *M. fortuitum* recovered from agar plates containing Levofloxacin (LVX) and Clarithromycin (CLR).

| Strains/Isolates | Source | MIC range (mg/L) |  |  |  |  |
| --- | --- | --- | --- | --- | --- | --- |
|  |  | GEPO | LVX | CLR | MEM | LZD |
| *M. fortuitum*<br>ATCC 6841 | Wild type | 2 | 0.06 | 1 | 2 | 2 |
| *M. fortuitum*<br>LVX-resistant | Isolated from LVX-containing agar plates inoculated with wild type | 2 | 4 | 0.5 | 4 | 2 |
| *M. fortuitum*<br>CLR-resistant | Isolated from CLR-containing agar plates inoculated with wild type | 2 | 0.12 | >64 | 2 | 2 |

GEPO: Gepotidacin, LVX- Levofloxacin, CLR- Clarithromycin, MEM: Meropenem, LZD- Linezolid

### *In vivo* efficacy of GEPO

Since GEPO exhibited potent activity against various mycobacterial pathogens, we proceeded to establish whether *in vitro* activity of GEPO translates *in vivo* as well. To test GEPO’s *in vivo* activity, we utilized the murine neutropenic *M. fortuitum* bacteremia model since it mimics clinical presentations of NTM infections (21) and the results are plotted in Fig 4A-C. As can be seen in Fig 4A-C, GEPO (10 mg/kg) significantly reduced bacterial counts in lungs (~2 log_10_ CFU/mL), spleen (~2 log_10_ CFU/mL) and kidney (~1.2 log_10_CFU/mL), which is comparable to reduction seen in lungs (~1.5 log_10_ CFU/mL), spleen (~2 log_10_ CFU/mL) and kidney (~1.2 log_10_ CFU/mL) of LVX (10 mg/kg) treated group while AMK (100 mg/kg) reduced bacterial counts in lungs (~1.6 log_10_ CFU/mL), spleen (~2 log_10_ CFU/mL) and kidney (~0.54 log_10_ CFU/mL) at 18 dpi. Taken together, GEPO exhibits potent *in vivo* activity as compared to AMK at 10-fold less concentration.

**Figure 4:**
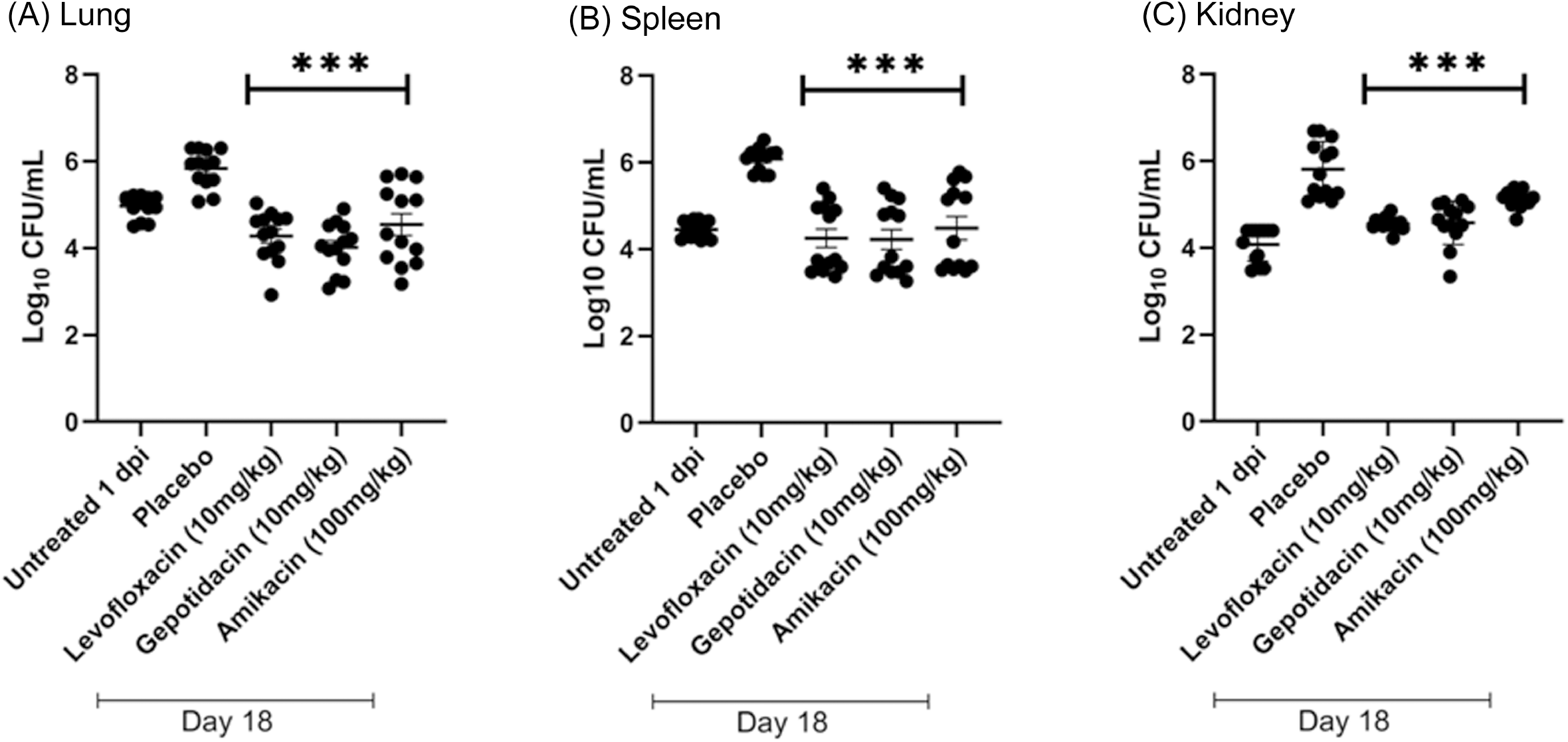
*In vivo* efficacy of Gepotidacin against *M. fortuitum* ATCC 6841. Bacterial load in various organs in murine bacteremia model. The bacterial load in various organs infected with a non-lethal dose of *M. fortuitum* ATCC 6841 (~5×10^6^ CFU) decreased significantly after treatment with Gepotidacin (10 mg/kg), LVX (10 mg/kg) and AMK (100 mg/kg). (**** equals <0.0001). All data were presented as mean ± s.d.

### Summary

Intrinsic and acquired drug resistance in mycobacteria is one of the factors responsible for a major increase in NTM infections observed worldwide (32) This is especially troubling since most of NTM infections occur in immunocompromised patients, thus negatively affecting their mortality(32). Similarly, the proportion of drug-resistant TB (DR-TB) continues to increase worldwide (1,6) despite the availability of an approved multi-drug regimen DOTS. Thus, identifying novel drugs active against mycobacterial pathogens which escape existing drug-resistance mechanisms is an urgent unmet medical need.

In this context, we have identified GEPO, a novel, first-in-class triazaacenapthylene molecule that inhibits bacterial type II topoisomerases by interacting with GyrA subunit of bacterial DNA gyrase and ParC subunit of bacterial topoisomerase IV through a unique mechanism to exhibit exquisite activity against various mycobacterial pathogens including those resistant to FQ. When tested in the time-kill assay, GEPO exhibits concentration-dependent bactericidal activity against both NTM and Mtb and synergizes with MER, MOX, CIP, LZD and AMK against NTM’s and INH while not interacting with RIF, ETB and STR against Mtb. GEPO exhibits potent intracellular infection clearing potential, against NTM’s, and shows a strong PAE of 12 h, which is comparable to AMK. Furthermore, *M. fortuitum* ATCC 6841 does not generate resistance to GEPO, while under the same conditions, it generates stable resistance to LVX. Additionally, when tested in the murine neutropenic bacteremia model, GEPO is more potent than AMK in reducing bacterial load in various organs at 10 fold lower concentration. Interestingly, Bulik *et al.,* have demonstrated that GEPO exhibits an *in vivo* PAE of ~3-12 h against S*. aureus*, which compares very well with the *in vitro* PAE reported by us (29, 30). Additionally, Bulik *et al.,* identified free-drug plasma AUC/MIC ratio as the PK-PD index most associated with *in vivo* efficacy of GEPO, with reported C_max_ of 4.5-14.3 mg/l in fasted and fed state respectively, which is very much in agreement with MIC against mycobacterial pathogens including FQ-resistant *M. abscessus* ATCC 19977 (29).

Taken together, GEPO is a broad-spectrum antibacterial agent with a novel mechanism of action that has demonstrated a safety profile consistent with other approved antibiotics in phase I clinical trials. PK-PD profiling of GEPO is further warranted for treating respiratory infections caused due to mycobacterial and other pathogens (25). These characteristics highlight GEPO’s stature as a promising, new antibacterial molecule targeting recalcitrant drug-resistant mycobacterial infections with a unique mode of action that is largely unaffected by existing resistance mechanisms as well as its inability to induce resistance.

## Acknowledgements

MNA and SS thank Council for Scientific and Industrial Research (CSIR),and TG thanks Department of Science and Technology (DST), Women Scientist Scheme (WOS-A) for their fellowships. The ORCID ID’s for MNA: https://orcid.org/0000-0002-9300-6107, TG: https://orcid.org/0000-0002-1997-6279, SS: https://orcid.org/0000-0003-0397-5498, SC: https://orcid.org/0000-0001-8823-6074 and ADG: https://orcid.org/0000-0001-9014-1904. This manuscript bears CDRI communication number XXXX.

## Ethics statement

The use of mice for infectious studies (IAEC/2019/114) was approved by Institutional Animal Ethics Committee at CSIR-CDRI, Lucknow.

## Transparency declarations

None to declare

